# Integrative Multi-Omics Analysis Identifies a Functional Enhancer Driving Tumorigenesis in Head and Neck Squamous Cell Carcinoma

**DOI:** 10.64898/2026.02.03.703438

**Authors:** Pengfei Xu, Xiaorong Liu, Jane Jingya Pu, Xiaoling Lu, Victor Lee, Evelyn Yilin Wang, Qianhui Jiang, Dan Yu, Bin Yan, Jinlin Song, Qi Zhong, Ping Yang, Nhan L. Tran, Xinyuan Guan, Yu-Xiong Su, Junwen Wang

## Abstract

Enhancer is a critical epigenetic feature in head and neck squamous cell carcinoma (HNSCC), yet the functional roles of individual enhancers remain poorly understood. Here, we conducted an integrative multi-omics analysis based on publicly available ATAC-seq, H3K27ac ChIP-seq, transcriptomic profiling, and genetic association datasets to systematically map HNSCC-associated enhancers and their target genes. Integrating ATAC-seq and H3K27ac ChIP-seq, we identified 20,362 enhancers and 18,040 enhancer-associated genes, highlighting widespread regulatory complexity. Functional characterization of the *TERT*-associated enhancer GH05J001312 (GRCh38/hg38: chr5:1312099-1317743) revealed strong transcriptional activity in HNSCC. CRISPR-mediated deletion of a core sequence significantly reduced *TERT* expression, impaired cellular proliferation *in vitro*, and suppressed tumor growth *in vivo*, confirming its role as a key cis-regulatory element. RNA-seq analysis of enhancer-edited cells uncovered 742 differentially expressed genes enriched in cancer-related pathways, including MAPK and IL-17 signaling, indicating a broad transcriptional impact. Collectively, our findings establish GH05J001312 as a functional enhancer driving oncogenic programs in HNSCC and suggest enhancer-targeted strategies as a potential therapeutic avenue.

## Introduction

Head and neck squamous cell carcinoma (HNSCC) is one of the most aggressive malignancies worldwide, characterized by extensive molecular heterogeneity and poor clinical outcomes despite advances in surgery, radiotherapy, and targeted therapies such as cetuximab ^1^. Overall survival has improved only marginally in recent decades, underscoring the need for novel therapeutic strategies ^2^.

Emerging evidence indicates that epigenetic dysregulation, particularly enhancer remodeling, plays a pivotal role in shaping oncogenic transcriptional programs that drive HNSCC initiation, progression, and therapeutic resistance ^3^. Enhancers are distal cis-regulatory elements that integrate chromatin accessibility, histone modifications, and transcription factor binding to orchestrate cell type-specific gene expression ^4-6^. Aberrant activation of oncogenic enhancers and enhancer-associated gene reprogramming has been recognized as a major contributor to tumorigenesis across multiple cancer types ^6^. Furthermore, perturbation of enhancers and enhancer-associated transcription factors selectively inhibits tumor growth and metastasis in preclinical models ^7^.

In HNSCC, enhancer-driven regulatory networks have been implicated in cancer stemness and progression. For example, a super-enhancer-mediated LIF/LIFR-STAT3-SOX2 feedback loop promotes stem-like properties in HNSCC ^3^, and *IGF2BP2* has been identified as a hub enhancer-associated gene exhibiting aberrant expression in HNSCC tissues ^8^. However, the enhancer landscape of HNSCC and the functional relevance of individual enhancers remain incompletely understood.

Multi-omics strategies have been applied to the identification of molecular biomarkers and regulatory pathways that serve as indicators of pathogenesis and prognostic factors in HNSCC ^9^. To systematically characterize enhancer-mediated transcriptional regulation in HNSCC, we performed an integrative multi-omics analysis combining ATAC-seq, H3K27ac ChIP-seq, transcriptomic profiling, and genetic association data. Through this approach, we identified a set of HNSCC-associated enhancers and prioritized their putative target genes. Notably, the enhancer GH05J001312 (GRCh38/hg38: chr5:1312099-1317743), together with its putative target gene TELOMERASE REVERSE TRANSCRIPTASE (*TERT*), emerged among the top candidates showing coordinated epigenomic and transcriptional alterations. *TERT*, the catalytic subunit of telomerase, functions as an oncogene through both telomere-dependent and telomere-independent mechanisms ^10^. While promoter mutations and methylation are the most widely discussed mechanisms of *TERT* regulation in HNSCC ^11,12^, the contribution of distal enhancers to *TERT* regulation has not been fully defined. Notably, the enhancer GH05J001312 has been implicated in *TERT* regulation in hematologic malignancies ^13^, but its role in HNSCC remains unknown. Here, we further functionally characterized GH05J001312 as a *TERT*-associated enhancer and demonstrated its critical role in promoting oncogenic transcriptional programs and tumor growth. Our findings provide new insights into enhancer-mediated regulation in HNSCC and highlight enhancer-targeted strategies as a potential therapeutic avenue.

## Results

### Genome-wide Identification of Enhancers and Associated Genes in HNSCC Using ATAC-seq

To delineate the enhancer landscape in HNSCC, we re-analyzed three ATAC-seq datasets to map genome-wide enhancer activity and elucidate their functional significance ^14^. An enrichment of ATAC-seq reads around transcription start sites (TSSs; ±3 kb) was observed across all datasets, which is a characteristic pattern expected for ATAC-seq (Figure 1A). Peak annotation analysis of HNSCC cell lines revealed that a large proportion of the peaks were located in promoters, introns, and distal intergenic regions (i.e., putative enhancer regions), with a notable enrichment in promoter regions (≤1 kb) (Figure 1B). We roughly defined these distal peaks as HNSCC-associated enhancers, yielding a total of 41,827 enhancers and 35,132 enhancer-associated genes (Figure 1C, Table S1-3). Moreover, we found many genes have been well established as oncogenes driving HNSCC development, such as *TP53, MYC* and *EGFR* (Figure 1D-F) ^15-17^. Significant peaks were enriched in the promoters, TSS, and intronic regions of these genes, suggesting that they may play important roles in the development of HNSCC.

**Figure 1.**
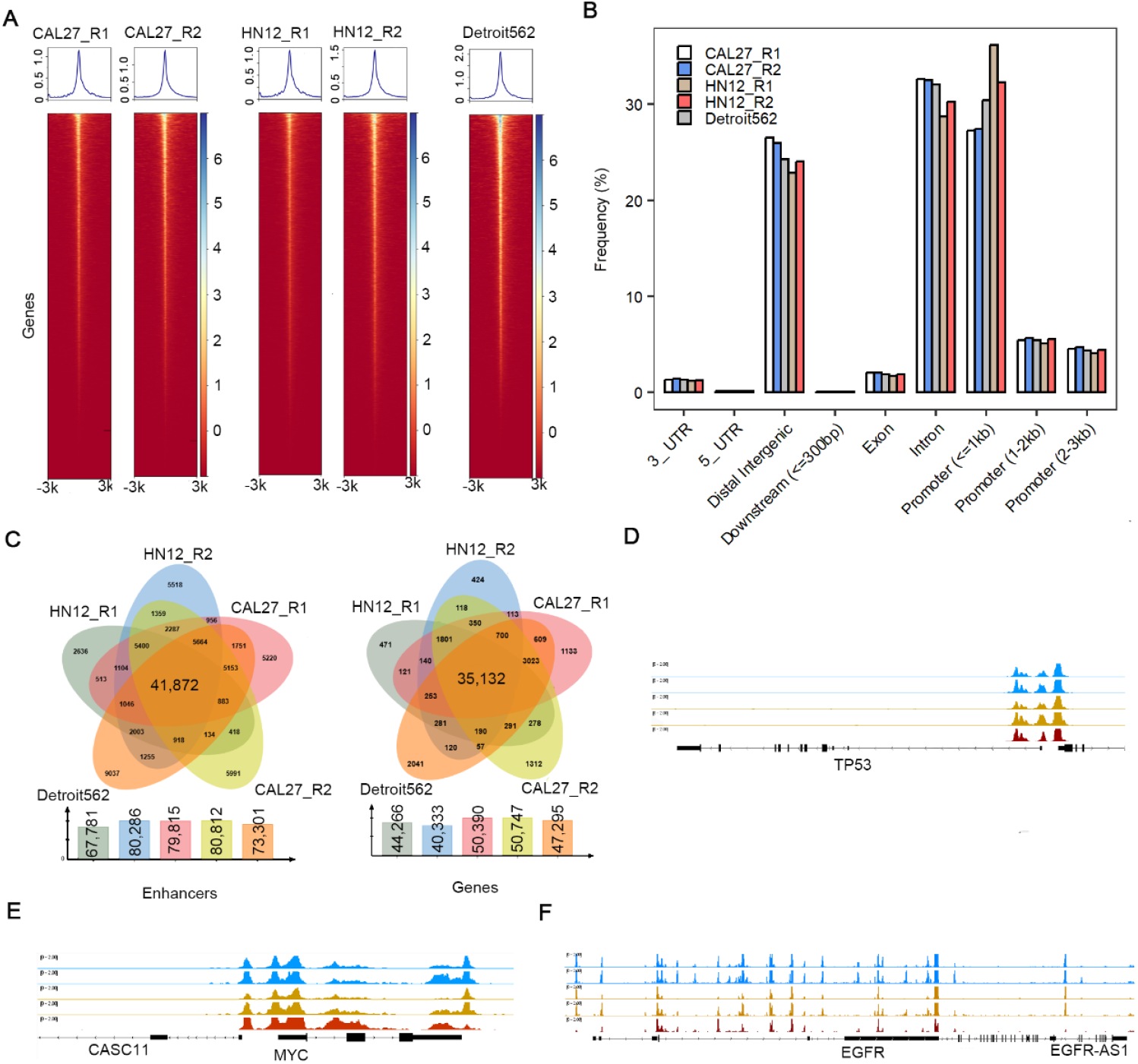
Genome-wide identifies of enhancer locations in 3 HNSCC cell lines. **(A)** Heatmaps displaying ATAC-seq signal enrichment at enhancer regions. **(B)** Genome-wide annotation of enhancer locations across three HNSCC cell lines. **(C)** Venn diagram showing common enhancers and putative enhancer target genes identified by ATAC-seq across three HNSCC cell lines. **(D-F)** ATAC-seq tracks showing signals in chromatin accessibility upstream of *TP53, MYC*, and *EGFR*.

### Integrative Epigenomic, Transcriptomic, and Genetic Data Identifies High Confidence Regulatory Drivers of HNSCC

To refine high-confidence regulatory elements, we re-analyzed a set of H3K27ac ChIP-seq dataset on the same three HNSCC cell lines used for ATAC-seq ^14^. Active enhancers were defined as H3K27ac peaks located within ±3 kb of transcription start sites. This approach allowed us to delineate the genome-wide enhancer landscape and characterize its biological roles. In total, we identified 45,972, 44,752, and 48,710 significant peaks from the CAL27, HN12, and Detroit562 cell lines, respectively. Among these, 81.00-85.44% were known enhancers, and 14.56-18.99% were newly identified in this study (Figure 2A, Table S4-6). An intersection analysis among the three datasets identified 21,953 enhancers and 18,502 enhancer-associated genes were shared (Figure 2B and C). furthermore, we found 20,362 enhancers and 18,040 genes were common identified between ATAC-seq and Chip-seq datasets (Figure 2D and E).

**Figure 2.**
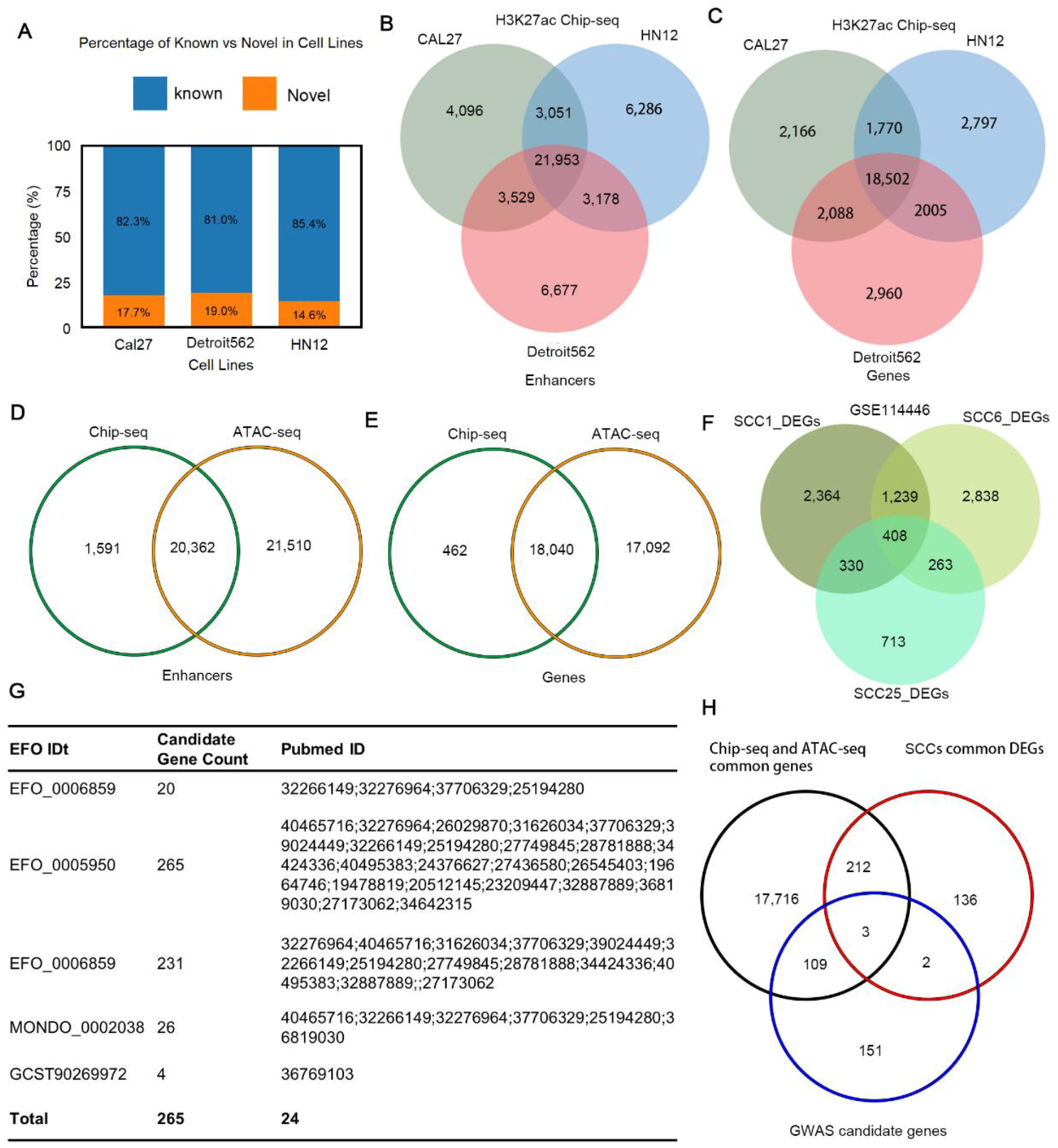
Integrative multi-omics analysis identifies candidate enhancers and their putative target genes in HNSCC. **(A)** The percentage of novel and known enhancers identified by H3k27ac chip-seq. **(B, C)** Venn diagrams showing common enhancers and putative enhancer target genes identified by H3K27ac ChIP-seq across three HNSCC cell lines. **(D, E)** Venn diagrams showing common enhancers and putative enhancer target genes identified by ATAC-seq and H3K27ac ChIP-seq. **(F)** RNA-seq identifies commonly DEGs) after JQ1 treatment across three HNSCC cell lines. **(G)** List of GWAS candidate genes associated with HNSCC. **(H)** Venn diagram showing common genes shared among ATAC-seq, H3K27ac ChIP-seq, and the GWAS catalog.

Transcriptional changes induced by cetuximab can be detected genome-wide shortly after treatment ^18^. Differential expression analysis of three HNSCC cell lines (SCC1, SCC6, and SCC25) revealed that exposure to this anti-EGFR therapy alters the transcriptional profiles of hundreds of genes ^18^. Intersection analysis of the three DEG datasets identified 408 genes that were consistently differentially expressed following cetuximab treatment compared with the mock control (Figure 2F). Furthermore, we queried the GWAS Catalog database (https://www.ebi.ac.uk/gwas/) and retrieved all candidate genes associated with HNSCC risk/development, yielding a total of 265 candidate genes (Figure 2G). Finally, by integrating the common genes identified from ATAC-seq and ChIP-seq with the DEGs from cetuximab treatment and the GWAS catalog, we found that three genes were shared across all datasets (Figure 2H). The three genes were *SLC7A7, TERT* and *SLCO4A1*. The three genes have previously been implicated in tumorigenesis and therapeutic responses across multiple cancer types ^19-21^, suggesting their potential functional relevance in HNSCC. *SLC7A7*, which encodes a cationic amino acid transporter, has been associated with metabolic reprogramming in cancer cells ^20^. *TERT*, a well-established oncogene, plays a central role in telomere maintenance and is frequently activated in HNSCC through promoter mutations ^21^. *SLCO4A1*, an organic anion transporter, may influence drug uptake and cellular homeostasis ^19^. The convergence of ATAC-seq, ChIP-seq, cetuximab-induced transcriptomic alterations, and genetic association evidence highlights these genes as high-confidence candidates for further mechanistic investigation. Collectively, our integrative analysis provides a refined set of regulatory and functional targets that may be associated with HNSCC susceptibility and therapeutic response.

### Expression and Clinical Significance of *SLC7A7, TERT*, and *SLCO4A1* in HNSCC Progression

To further evaluate the roles of *SLC7A7, TERT* and *SLCO4A1* in the development of HNSCC, we first analyzed the expression patterns of these three genes in a public HNSCC cohort ^13^. Compared with adjacent normal tissues, all three genes exhibited significant expression changes between tumor and normal samples. *SLC7A7* and *TERT* were markedly upregulated in tumors, whereas *SLCO4A1* showed lower expression in cancer tissues (Figure 3A). We also generated a cohort consisting of 12 patients and performed bulk RNA-seq on paired tumor and adjacent normal tissues. In our dataset, the expression patterns of *SLC7A7* and *TERT* were consistent with those observed in public dataset, suggesting that these two genes may play important roles in the pathogenesis of HNSCC (Figure 3B). Further, we examined whether these two genes influence HNSCC progression. Using clinical data downloaded from the TCGA database, we analyzed the expression levels of these two genes across different tumor stages. We found that the expression of these two genes changes during HNSCC progression. Specifically, the expression levels of *SLC7A7* and *TERT* both increase significantly from tumor stage I to tumor stage IV (Figure 3C and E). These results suggested that these two genes serve as a valuable predictive biomarker and potential therapeutic target in HNSCC ^22^. Moreover, two enhancers, GH14J022817 and GH05J001312, were annotated upstream of *SLC7A7* and *TERT*, respectively. Notably, cetuximab altered the accumulation of reads covering these two annotated enhancer regions, concomitant with the changed expression levels of *SLC7A7* and *TERT*, suggesting that these genes may be regulated by their corresponding enhancers (Figure 3D and 3F).

**Figure 3.**
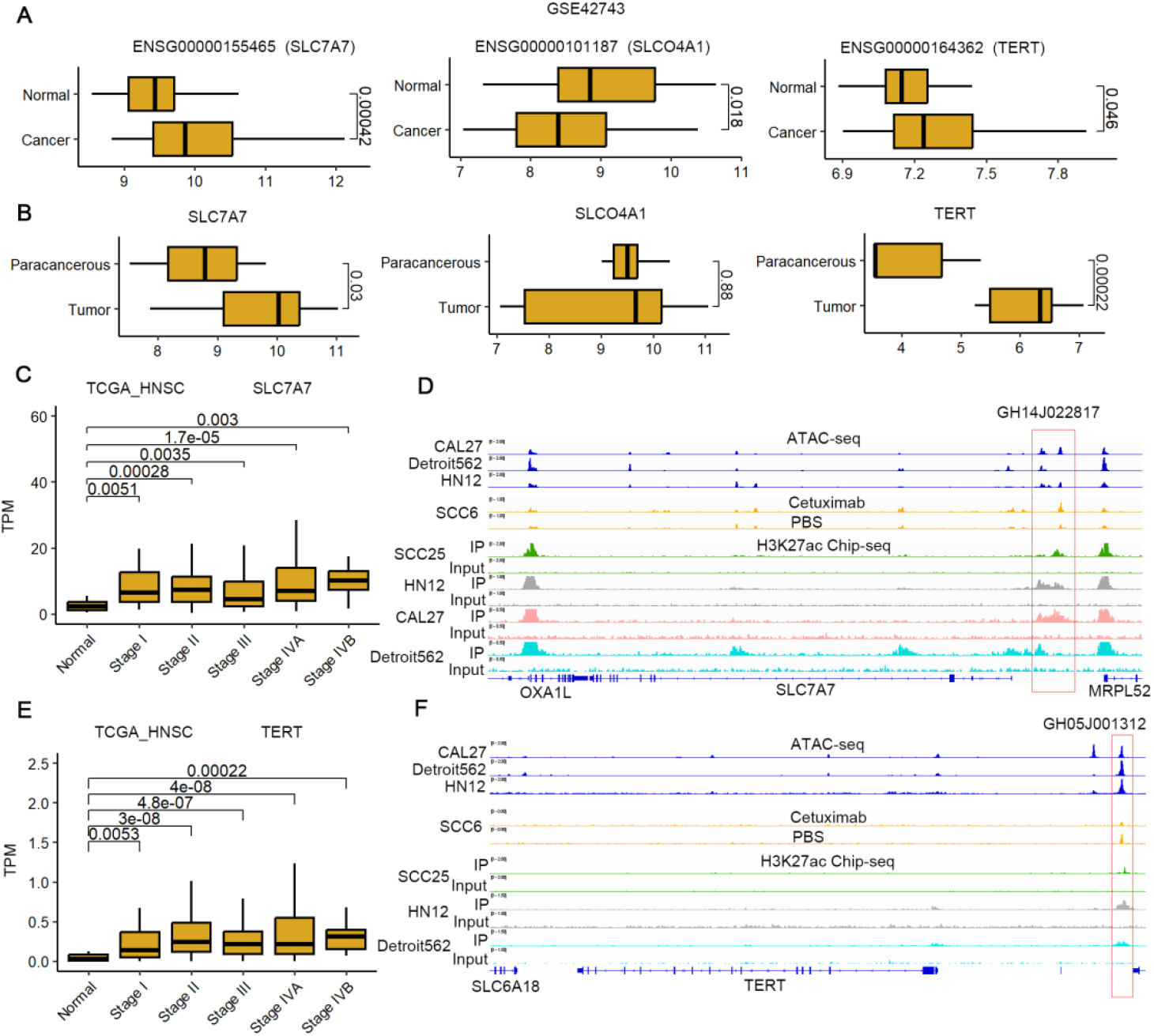
Expression and clinical significance of 3 genes HNSCC progression. **(A, B)** The average expression of *SLC7A7, SLCO4A1* and *TERT* in a public HNSCC dataset (A) and private HNSCC dataset (B). The pvalue was tested by the Wilcoxon test. **(C, E)** The clinical significance of *SLC7A7* (C) and *TERT* (E) in TCGA-HNSCC. The pvalue was tested by the Wilcoxon test. **(D, F)** ATAC-seq and H3k27ac ChIP-seq tracks showing signals upstream of *SLC7A7* (D) and *TERT* (F).

### A Core Sequence Within the GH05J001312 Enhancer Promotes TERT Transcription and Tumor Growth in HNSCC

In cancer, TERT stabilization leads to constitutive self-renewal preventing DNA damage response. However, the functional of its enhancer hasn’t been well understood in HNSCC. Based on the analysis of peak positions in ChIP-seq and ATAC-seq datasets, a candidate core DNA sequence of this enhancer element was identified (Table S7) and corned into a luciferase reporter system. As illustrated in Figure 4A, this enhancer consistently induced much higher luciferase activities in two HNSCC cell lines as compared a promoter only. This result indicates that this enhancer is functionally active in HNSCC and potentially serving as a key regulatory element driving its target gene expression in this cancer type. To further examinate the function of this core DNA sequence, we performed CRISPR-KO to knock out this core DNA sequence from this enhancer. Three mutant cell lines were obtained after validated by Sanger sequencing (Figure 4B). The expression level of *TERT* was quantified by qPCR, the result showed that the expression level of *TERT* in the three was decreased as compared to control CAL27 cell lines (Figure 4C). Taken together, these results demonstrate that the identified enhancer core sequence is required for maintaining *TERT* expression in HNSCC, supporting GH05J001312 as a functional enhancer in *TERT* transcriptional regulation.

**Figure 4.**
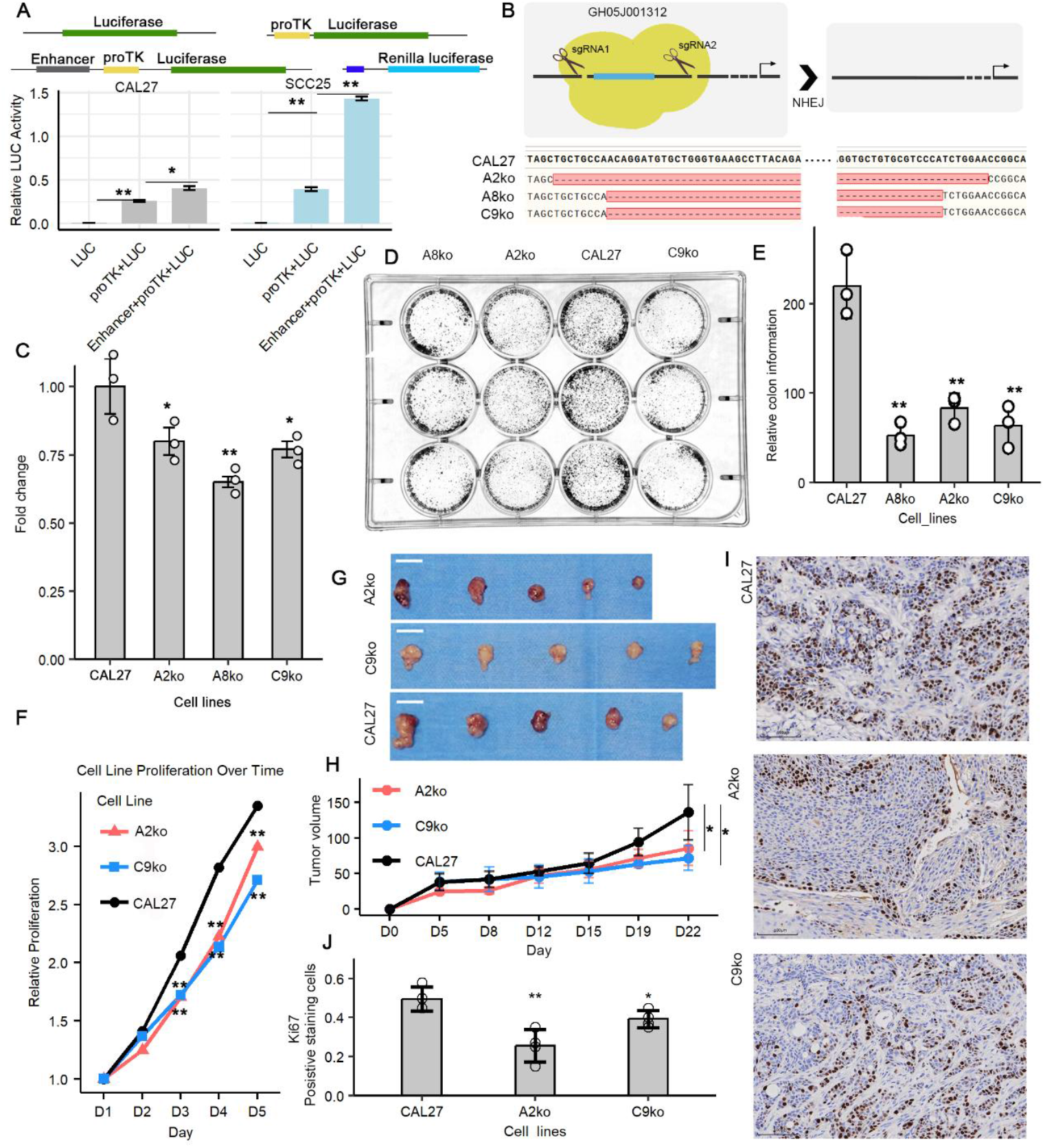
Deletion of a core fragment within enhancer GH05J001312 suppresses tumor growth by downregulating *TERT* expression. **(A)** The core enhancer region was cloned upstream of the luciferase promoter and further used for luciferase reporter assays (n = 3; student’s t-test), *p < 0.05; **p < 0.01. **(B)** A core fragment within enhancer GH05J001312 was deleted in three independent cell lines. **(C)** Deletion of a core fragment within enhancer GH05J001312 suppresses *TERT* expression (n = 3; student’s t-test), Data are shown in means ± SE. *p < 0.05; **p < 0.01. **(D, E)** Colony formation assays showed impaired colony forming ability in mutant cells compared with controls (n = 3; student’s t-test), *p < 0.05; **p < 0.01. **(F)** The cell proliferation of mutant cells and control cells (n = 3; student’s t-test). **(G, H)** Tumor growth in Balb/c nude mice. Tumors in the control and mutant groups were collected at the end of the experiments. Tumor growth was assessed by tumor volume **(H, J)** (n = 5, student’s t-test). *p < 0.05; **p < 0.01. (I) Representative images **(I)** and quantitative analysis **(J)** of Ki67 expression in control and mutant cell lines (×200, scale bars, 100 μm) (n = 5, student’s t-test).

Cellular growth and colony formation assays were subsequently performed to evaluate the effects of manipulating this enhancer on the proliferative capacity of HNSCC cells. We found that knocking out this core DNA sequence decreased the proliferation capacity of HNSCC cells (Figure 4D-F). To determine the contribution of GH05J001312 to HNSCC tumor cell growth in vivo, CAL27 and mutant cell lines were inoculated into nude mice. Consistent with our in vitro findings, we observed that the deletion significantly suppressed tumor growth, as reflected by reduced tumor volume (Figure 4G and H). Furthermore, mice implanted with the mutant cell lines exhibited a higher body weight compared to those implanted with CAL27 cells, although this difference did not reach statistical significance (Figure S1). Ki-67 is a nuclear protein commonly used as a marker of tumor cell proliferation ^23^. Subsequently, IHC stains for xenograft tissue extracted from nude mice demonstrated that knocking out core DNA sequence can markedly downregulate Ki-67, which is related to cancer growth (Figure 4I and J).

Taken together, these results identify GH05J001312 as a functional enhancer that drives *TERT* expression and contributes to the proliferative and potentially tumorigenic capacity of HNSCC cells. This core DNA sequence therefore represents a promising candidate for future therapeutic strategies targeting TERT-dependent tumor maintenance in head and neck cancer.

### Transcriptomic Profiling and Functional Enrichment Analysis Following GH05J001312 Enhancer Mutation

Bulk RNA sequencing (RNA-seq) was performed to compare the gene-expression profiles of HNSCC cells with or without GH05J001312 mutation, aiming to investigate the potential mechanisms underlying the function of this enhancer. RNA-seq libraries generated 21-24 million paired-end reads per sample, of which ∼95% were uniquely mapped to the human reference genome (Table 1). Gene expression was quantified and normalized to Transcripts Per Million (TPM). Principal component analysis (PCA) of the normalized expression data demonstrated that biological replicates from the same cell line clustered closely together, indicating strong technical and biological reproducibility (Figure 5A). The correlation heatmap analysis shows extremely high correlations between cell lines (Cor > 0.98), indicating that knocking out a core DNA sequence only caused a minor perturbation to the CAL27 genome (Figure 5B). To evaluate differences in gene expression between the CAL27 cell line and the mutant cell lines, we performed differential expression analysis using the criteria |log_2_FoldChange| ≥ 0.58 and p-value < 0.05. The results revealed that 3,666 genes were significantly differentially expressed in CAL27 vs. A2ko (2,344 upregulated and 1,322 downregulated in A2ko relative to CAL27). Similarly, 2,811 genes were differentially expressed in CAL27 vs. A8ko (979 upregulated and 1,832 downregulated in A8ko), and 2,974 genes were differentially expressed in CAL27 vs. C9ko (1,578 upregulated and 1,396 downregulated in C9ko) (Figure 5C). Intersection analysis among the three clusters identified 742 genes that were shared (Figure 5D). This result indicates that editing an enhancer can influence the expression of hundreds of genes. Therefore, studying enhancer-regulated genes is a potential approach for contributing to HNSCC therapy. To explore the function of these 742 genes, we performed Gene Ontology (GO) enrichment analysis. The results showed that they were significantly enriched for terms related to multicellular organismal processes, developmental processes, anatomical structure development, and cell differentiation (Figure 5E). Kyoto Encyclopedia of Genes and Genomes (KEGG) analysis of these 742 genes revealed their significant enrichment in several pathways, including pathways in cancer, the MAPK signaling pathway, the cytoskeleton in muscle cells, and cytokine-cytokine receptor interaction (Figure 5F). Taken together, these results demonstrate that knocking out the core DNA sequence in an HNSCC cell line leads to widespread alterations in gene expression.

**Table 1.** Summary of Bulk RNA-Seq Data.

**Figure 5.**
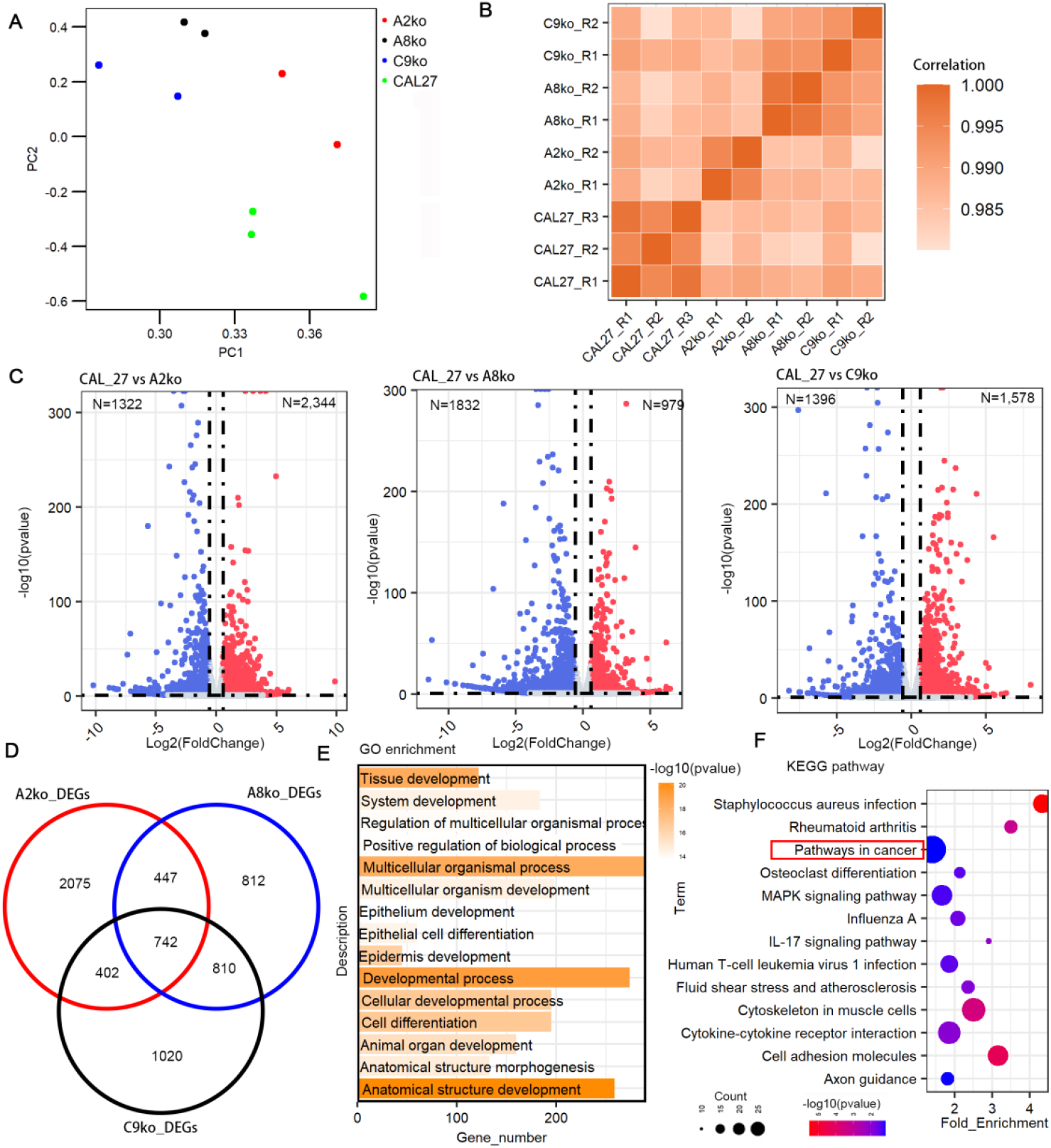
Global view of the impacts of knocking out the functional enhancer using RNA-seq. **(A)** PCA of normalized RNA-seq data. **(B)** Heatmap showing the correlation among the RNA-seq libraries. **(C)** Volcano plot of DEGs between control and mutant cell lines. **(D)** Venn diagram showing the overlap of DEGs among the three libraries. **(E)** GO enrichment analysis of the DEGs. **(F)** KEGG enrichment analysis of the DEGs.

### qPCR Validation of the Identified Differentially Expressed Genes

In our study, we found that DGEs were enriched in “Pathways in cancer”, “MAPK signaling pathway”, and “IL-17 signaling pathway”. These pathways have been reported to participate in various aspects of cancer biology, including tumor initiation, proliferation, survival, and immune regulation ^24,25^. Their enrichment suggests that the identified DGEs may contribute to malignant progression through dysregulation of key oncogenic and inflammatory signaling networks, thereby providing potential mechanistic insights and therapeutic targets for further investigation. To verify the reliability of the RNA-seq data, quantitative PCR (qPCR) was performed to confirm the expression changes of the selected genes. Those genes that exhibited similar expression changes across the three mutant cell lines were selected for further examination of their expression levels. For instance, *FGFR3, CCNA1, PGF* and *ETS1*, which were enriched in “Pathways in cancer” and have been reported to play critical roles in cancer ^26-29^. *MAPK8IP2* and *TGFB2* were enriched in MAPK signaling pathway. All of these genes were differentially expressed between the control and mutant groups (Figure 6A). Consistent with the RNA-seq data, qPCR analysis confirmed the expression changes of these genes in the mutant cell lines compared with controls. Collectively, these results suggest that GH05J001312 is associated with widespread alterations in cancer-related gene expression programs.

**Figure 6.**
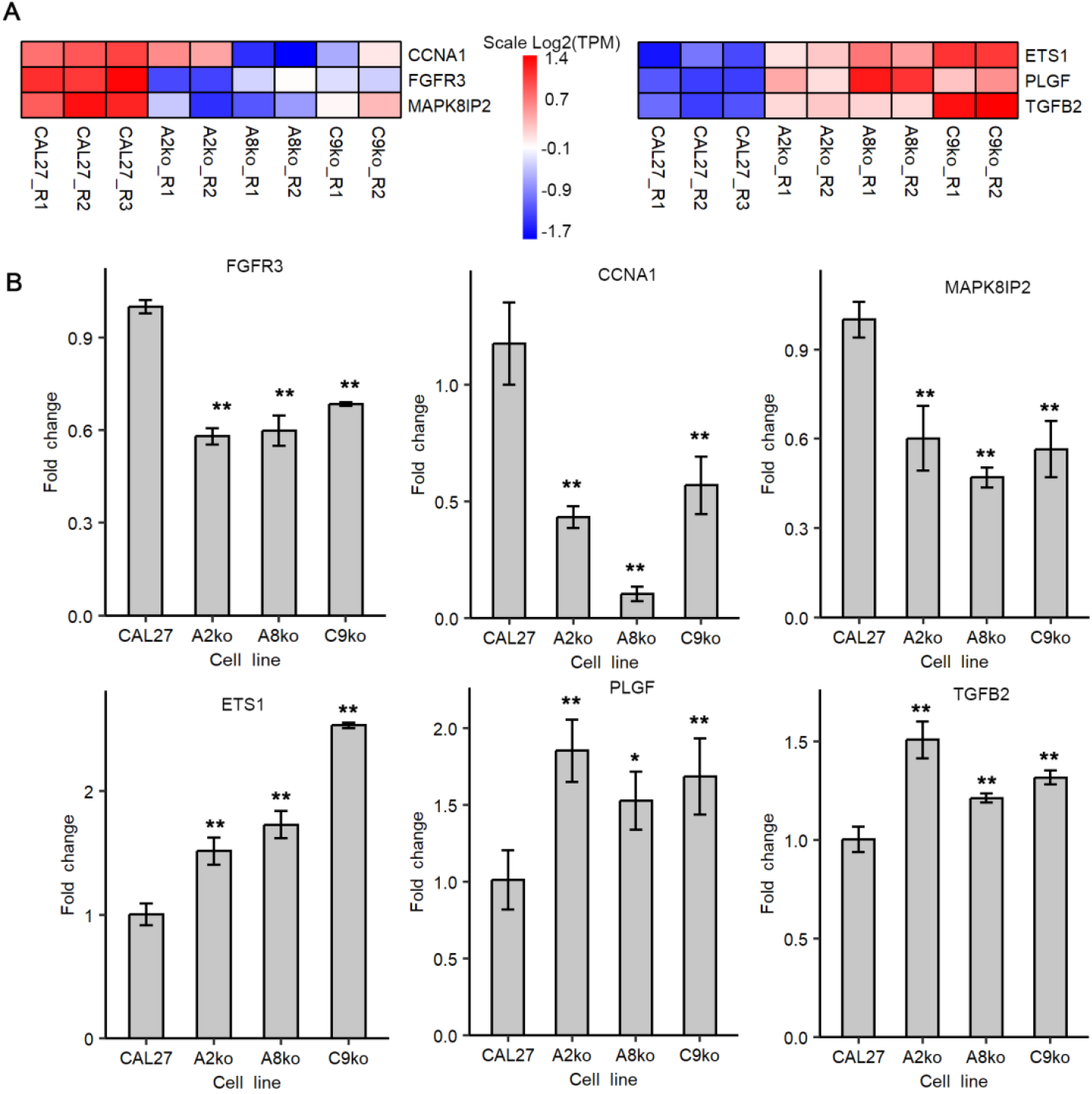
qPCR confirms the expression changes of DEGs identified by RNA-seq. (n = 3; student’s t-test), Data are shown in means ± SE. *p < 0.05; **p < 0.01.

## Discussion

Over the past years, tremendous progress has been made in elucidating the transcriptional deregulation that underlies tumorigenesis. This progress has largely been driven by the identification of key regulatory elements, both trans-acting factors and cis-acting elements, and the characterization of their mechanisms of action ^30^. Aberrant activation of oncogenic enhancers and super-enhancers has been demonstrated to be an important contributor to tumorigenesis across multiple cancer types ^31,32^ Our study identifies GH05J001312 as a functional enhancer that activates *TERT*, promoting malignant progression in HNSCC. These findings underscore the critical role of enhancer-mediated regulation in cancer biology and highlight enhancers as promising prognostic markers and therapeutic targets.

In this study, we systematically integrated epigenomic, transcriptomic, and genetic datasets to uncover key regulatory elements and target genes involved in HNSCC development. By combining ATAC-seq, H3K27ac ChIP-seq, GWAS data, and cetuximab-induced transcriptional changes, we constructed a comprehensive enhancer atlas and identified high-confidence regulatory elements and genes associated with HNSCC (Figure 1 and 2). Among these, *SLC7A7, TERT*, and *SLCO4A1* emerged as robust candidates involved in HNSCC development and therapeutic response. Expression profiling in public and in-house cohorts revealed significant dysregulation of these genes in tumors and progressive upregulation of *SLC7A7* and *TERT* during tumor advancement (Figure 3). Among these, the enhancer GH05J001312 emerged as a particularly strong candidate, supported by converging evidence from chromatin accessibility, enhancer-associated histone modifications, and genetic signals (Figure 3F).

To further elucidate the function of GH05J001312 in HNSCC, we first used luciferase reporter assays to assess the transcriptional activation driven by its core enhancer sequence. We observed that this core sequence strongly enhanced *LUC* transcription in HNSCC cells, confirming that GH05J001312 functions as an active cis-regulatory element (Figure 4A). Functionally, we demonstrated that GH05J001312 acts as a TERT-activating enhancer in HNSCC, as CRISPR-mediated mutation of this element markedly reduced *TERT* expression, impaired cell proliferation, and suppressed tumor growth both in vitro and in vivo (Figure 4B-J). These findings not only establish GH05J001312 as a functional regulator of *TERT* but also provide mechanistic insights into how enhancer regulation contributes to the aggressive growth phenotype characteristic of HNSCC. Although *TERT* activation in cancer has typically been attributed to promoter mutations, our study highlights enhancer-mediated regulation as an additional and potentially therapeutically relevant layer of transcriptional control.

Transcriptome profiling demonstrated that GH05J001312 deletion led to broad alterations in gene expression, with hundreds of genes being differentially expressed across all mutant cell lines. Notably, the commonly dysregulated genes were enriched in pathways closely associated with cancer biology, including “Pathways in cancer,” “MAPK signaling pathway”, “IL-17 signaling pathway”, and cytokine-receptor interactions. The involvement of these pathways suggests that GH05J001312 deletion disrupts not only *TERT* transcription but also a wider regulatory network influencing oncogenic signaling, cell differentiation, and tumor-microenvironment interactions (Figure 5). The consistency of expression changes across three independent mutant cell lines further strengthens the conclusion that GH05J001312 has a reproducible and widespread regulatory impact.

qPCR analyses of selected genes, including *FGFR3, CCNA1, PGF, ETS1, MAPK8IP2* and *TGFB2* validated the RNA-seq findings. These genes are known drivers of cancer progression and are implicated in proliferation, angiogenesis, cell-cycle regulation, and transcriptional control, reinforcing the functional significance of the affected pathways (Figure 6). Collectively, these results indicate that GH05J001312 plays a broader biological role beyond regulating *TERT* alone and may influence multiple oncogenic processes in HNSCC.

Taken together, our findings identify GH05J001312 as a previously uncharacterized enhancer with substantial impact on cancer-related transcriptional programs in HNSCC. This enhancer not only drives *TERT* expression but also modulates a large network of downstream genes involved in tumor growth and malignant progression. These results underscore the importance of non-coding regulatory elements in cancer development and highlight GH05J001312 as a promising molecular target for therapeutic intervention. Future studies dissecting the transcription factors and chromatin interactions governing GH05J001312 activity will further clarify its regulatory mechanisms and may uncover additional opportunities for targeted therapy in head and neck cancer.

## Methods and materials

### Cell Lines and Cell Culture

Human head and neck squamous cell carcinoma (HNSCC) cell lines CAL27 (cat: CRL-2095) and SCC25 (cat: CRL-1628) were obtained from the American Type Culture Collection (ATCC). Cells were maintained in Dulbecco’s Modified Eagle Medium (DMEM; Gibco) supplemented with 10% fetal bovine serum (FBS; Gibco) and 1% penicillin/streptomycin, and cultured at 37 °C with 5% CO_2_.

### ATAC-seq and H3K27ac ChIP-seq Data Processing and Enhancer Identification

Publicly available ATAC-seq and H3K27ac ChIP-seq datasets from three HNSCC cell lines were downloaded from the GEO database (GSE128275). The adaptors were trimmed and then aligned to the human reference genome (hg38) using Bowtie2 (Version 2.5.4). Peaks were called by MACS3 with default parameters (Version 3.0.4). Peaks located in promoter, intronic, and distal intergenic regions were annotated using ChIPseeker R package and defined as putative enhancers. Peaks located ±3 kb of transcription start site (TSS) were defined as active enhancers. Enhancer-associated genes were assigned based on the nearest TSS.

### Integration of ATAC-seq, ChIP-seq, DEGs, and GWAS Data

Differentially expressed genes (DEGs) induced by cetuximab treatment in SCC1, SCC6, and SCC25 cell lines were obtained from previously published datasets ^18^. Candidate HNSCC-associated genes were retrieved from the GWAS Catalog. Intersections among enhancer-associated genes, DEGs, and GWAS candidates were performed using the JVeen online website (https://jvenn.toulouse.inra.fr/app/example.html) to identify high-confidence regulatory target genes.

### RNA Extraction and RNA-seq Analysis

Total RNA from CAL27 and GH05J001312-KO cells was extracted using TRIzol (Invitrogen). RNA integrity was assessed with an Agilent 2100 Bioanalyzer. Poly(A)-selected RNA libraries were constructed using the Illumina TruSeq Stranded mRNA Library Preparation Kit and sequenced on the NovaSeq platform. Reads were aligned to hg38 using STAR (Version STAR_2.7.11b), and gene level quantification was performed using featureCounts. Expression levels were normalized to TPM. DEGs were identified using DESeq2 with thresholds |log_2_FoldChange| ≥ 0.58 and p < 0.05. PCA and correlation analyses were performed using the ggpubr and pheatmap R packages.

### Gene Ontology and KEGG Pathway Analysis

The Gene Ontology enrichment of DEGs was conducted using the online website PANTHER (https://geneontology.org/). KEGG pathway enrichment analysis was performed using the DAVID Bioinformatics Resources (https://david.ncifcrf.gov/). A list of differentially expressed genes (DEGs) was uploaded using official gene symbols as identifiers. GO terms and KEGG pathways with adjusted p < 0.05 were considered significantly enriched.

### Construction of Enhancer Reporter Plasmid and Luciferase Assay

A candidate core sequence of the GH05J001312 enhancer was synthesized and cloned upstream of a minimal promoter in the PGL3-BASIC-TK luciferase reporter vector. CAL27 and SCC25 cells were transfected using Lipofectamine 3000 (Invitrogen). Luciferase activity was measured 48 h post-transfection using the luciferase activity was measured with the Luciferase Reporter Gene Assay Kit and Dual-Lumi™ Luciferase Assay Kit (A1222, Promega), with Renilla luciferase as an internal control.

### CRISPR-Cas9 system Mediated Enhancer Knockout

The core enhancer region of GH05J001312 was deleted using modified CRISPR-Cas9. In brief, Cas9 protein and gRNA were obtained through in vitro synthesis. The Cas9 protein and gRNA were electroporated into CAL27 cells. After 48 hours of electroporation, cells were collected and then sorted using a BD Influx cell sorter (BD Biosciences) and seeded into 96-well plates to initiate single-cell cloning. After PCR amplification and sequencing validation, three positive clones were obtained. Three confirmed knockout clones (A2ko, A8ko, and C9ko) were used for downstream experiments.

### Quantitative PCR (qPCR)

A total of 2□μg of high-quality total RNA (A260/280□=□2.00) was reverse-transcribed into cDNA using the PrimeScript™ RT Reagent Kit with gDNA Eraser (Takara, Otsu, Japan; Code No. RR047A). qPCR was performed using TB Green® Premix Ex Taq™ II (Tli RNase H Plus) (Takara, Otsu, Japan; Code No. RR820A) on StepOnePlus Real-Time PCR system (Applied Biosystems, Thermo Fisher Scientific, USA). Gene expression was normalized to GAPDH using the 2^−ΔΔCt^ method. Genes exhibiting consistent expression changes across all three mutant cell lines were selected for validation, the specific primers (Table S8) were download from OriGene (https://www.origene.com/services/custom-cloning).

### Cell Proliferation Assay and Colony Formation Assay

Cell proliferation was measured using the CCK8 (CCK8, Dojindo, Japan) assay. CAL27 and enhancer-knockout cells were seeded into 96-well plates. Approximately 1000-3000 cells/well were seeded in the 96-well plates. 10 ul agent was added at indicated time points and incubated at 37°C for another 2 h. Absorbance at 450 nm was measured using a microplate reader.

Cells (3000 per well) were seeded in 12-well plates and cultured for 10 days, with medium refreshed every 3 days. Colonies were fixed with 4% paraformaldehyde for 15 minutes and stained with 0.1% crystal violet. After washing and air drying, the colonies were further visualized and counted.

### HNSCC Xenograft Tumor Model and Immunohistochemistry (IHC)

All animal experiments were approved by the institutional animal care committee of Shenzhen Lingfu Topu Biotechnology Co., Ltd and performed in accordance with the National Institutes of Health Guide for the Care and Use of Laboratory Animals. BALB/c nude mice (4 weeks old) were subcutaneously injected with 1 × 10□ CAL27 or GH05J001312-KO cells. Tumor volume was measured every 3-4 days using the formula: Volume=1/2×((the longest diameter)×(the shortest diameter)^2^). Tumor tissues were harvested for endpoint analysis and immunohistochemistry.

IHC staining was performed as previously described ^33^. In brief, paraffin embedded xenograft tissues were sectioned at 5 μm. Slides were subjected to antigen retrieval followed by incubation with Ki-67 antibody (Abcam™; #ab15580; 1:400 dilution) primary antibody overnight at 4°C and secondary antibody (Abcam; #ab205718; 1:1000 dilution) for 1 h at room temperature. Tissue segments were finally subjected to visualization with a diaminobenzidine (DAB) Kit (Invitrogen) and imaging under a phase-contrast microscope (Leica™, Cat. #DMI 1).

## Acknowledgements

The work was funded by the oral health research and innovation fund of the faculty of dentistry (006010309) from the University of Hong Kong to P XU, and this research is funded by the Hong Kong Scholars Program. The work was funded by collaborative research fund of Hong Kong RGC (C7015-23G), seed funding for collaborative research (2207101590) from the University of Hong Kong to J Wang.

## Author contributions

PX and JW designed the study. PX and DY compiled all sequencing data and carried out the computational analysis with input from JW. PX and XL performed the molecular and biochemical experiments with input from JW. PX, JP, XL, QJ, XG, and YS collected the clinical samples. PX, BY and JW wrote the manuscript. VL, YW, JS, QZ, PY and NT provide valuable comments on this project. All authors read and approved the manuscript.

## Conflict of interest

The authors declare no conflict of interest.

## Data access

All the raw data and processed sequencing data generated in this study have been submitted to the NCBI under accession number of GSE128275 (ATAC-seq and H3K27ac chip-seq data). The enhancer list was acquired at the following URL: https://www.genecards.org/Download/File?file=GeneHancer_v5.25.bed. The transcriptome data reported in this study have been deposited in NCBI under accession number GSE313806.

## Ethics statement

This study was approved by the Institutional Review Board of the University of Hong Kong/Hospital Authority Hong Kong West Cluster (IRB Reference Number: UW 23-590). Informed consent was obtained from all patients for the research use of their tissues.

## Supporting information

The following materials are available in the online version of this article.

Table S1. Candidate enhancers identified in the CAL27 cell line using ATAC-seq.

Table S2. Candidate enhancers identified in the HN12 cell line using ATAC-seq.

Table S3. Candidate enhancers identified in the Detroit562 cell line using ATAC-seq.

Table S4. Candidate enhancers identified in the CAL27 cell line using H3K27ac chip-seq.

Table S5. Candidate enhancers identified in the HN12 cell line using H3K27ac chip-seq.

Table S6. Candidate enhancers identified in the Detroit562 cell line using H3K27ac chip-seq.

Table S7 The candidate core DNA sequence of GH05J001312 enhancer.

Table S8 The specific primers used in this study.

Figure S1 Mouse body weights after injection of HNSCC cell lines.

## Notes

### Competing Interest Statement

The authors have declared no competing interest.

