## Supplementary figures and images for "Integrative Multi-Omics Analysis Identifies a Functional Enhancer Driving Tumorigenesis in Head and Neck Squamous Cell Carcinoma"

### Figure S1

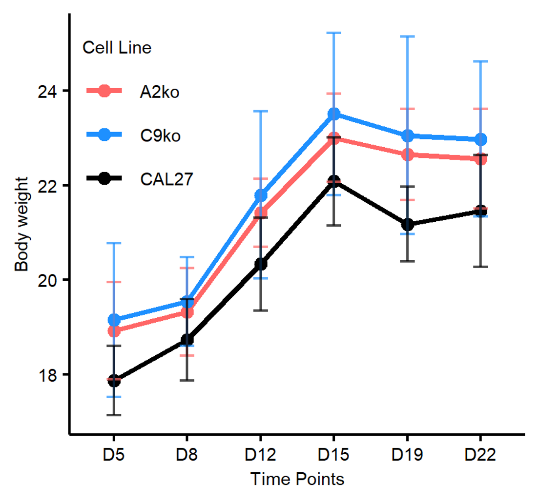

**Figure S1** Mice weights after injection of HNSCC cell lines. Data points represent mean values.
